# A newly identified three-domain C-type lectin associated with blood feeding in the tick *Ixodes ricinus*

**DOI:** 10.64898/2026.01.07.698205

**Authors:** Kateryna Kotsarenko, Ondřej Hajdušek, Martina Rievajová, Kateřina Bezděková, Pavlína Věchtová, Lenka Malinovská, Eva Paulenová, Josef Houser, Ján Štěrba, Libor Grubhoffer, Michaela Wimmerová

**Affiliations:** Central European Institute of Technology, Masaryk University, Kamenice 5, Brno, 62500, Czech Republic; National Centre for Biomolecular Research, Faculty of Science, Masaryk University, Kotlarska 2, Brno, 61137, Czech Republic; Faculty of Science, University of South Bohemia, Branisovska 1760, Ceske Budejovice, 37005, Czech Republic; Institute of Parasitology, Biology Centre of the Czech Academy of Sciences, Branisovska 31, Ceske Budejovice, 37005, Czech Republic; Department of Biochemistry, Faculty of Science, Masaryk University, Kotlarska 2, Brno, 61137, Czech Republic

**Author notes:** E-mails: Kateryna Kotsarenko –, Ondřej Hajdušek –, Martina Rievajová –, Kateřina Bezděková –, Pavlína Věchtová –, Lenka Malinovská –, Eva Paulenová –, Josef Houser –, Ján Štěrba –, Libor Grubhoffer –.

**Keywords:** C-type lectin, glycan array, innate immunity, blood feeding, *Ixodes ricinus*

## Abstract

*Ixodes ricinus* ticks are widely distributed throughout Europe and represent major vectors of tick-borne encephalitis virus and the Lyme borreliosis agent *Borrelia burgdorferi* sensu lato. In invertebrates, C-type lectins are commonly associated with innate immune functions, and several such lectins have been predicted in *I. ricinus*. Given the limited knowledge of lectin function in ticks, we characterized three carbohydrate-recognition domains (CRDs) of a novel C-type lectin identified in the *I. ricinus* transcriptome (IrCLec).

The tertiary structures of CRD1, CRD2, and CRD3, predicted using the AlphaFold 3 program, corresponded to the typical structure of C-type lectins. Conserved carbohydrate-binding motifs were identified in CRD3, whereas non-canonical motifs were present in CRD1 and CRD2. Recombinant His-tagged CRDs were produced and analysed for carbohydrate-binding activity. Glycan array analysis revealed binding of all three domains to selected glycans, while hemagglutination assays demonstrated pronounced binding activity of CRD1 and CRD2 toward human erythrocyte antigens of blood groups A, B, and O.

IrCLec expression was highest in the tick midgut and also detected in hemocytes, with expression levels increasing after blood feeding. RNAi-mediated silencing of IrCLec impaired blood feeding efficiency in tick nymphs. Together, these results indicate that IrCLec plays an important role in blood feeding and may additionally participate in lectin-mediated host-pathogen or host-blood component interactions.

## Introduction

*Ixodes ricinus* is the most important tick vector in Europe, transmitting the tick-borne encephalitis (TBE) virus and the Lyme borreliosis-causing spirochete *Borrelia burgdorferi* sensu lato (Grubhoffer et al., 2004). Although its vector competence is suggested to be closely linked to its immune system, knowledge about the immunity of *I. ricinus* remains limited compared to that of other arthropods. (Hajdušek et al., 2013; Kopáček et al., 2010; Taylor, 2006).

To date, two groups of proteins have been identified as antimicrobial agents in *I. ricinus*: defensins (Chrudimská et al., 2011, Tonk et al., 2014; Tonk et al., 2015;) and ixoderins (Honig Mondekova et al., 2017, Rego et al., 2005;). Moreover, the presence of C-type lectins (CTLs) has been suggested based on the observed Ca^2+^-dependent hemagglutinating activity in hemolymph (85 kDa protein), midgut (37, 60, 65, and 70 kDa proteins), and salivary glands (70 kDa protein) (Grubhoffer and Jindrák, 2013; Grubhoffer et al., 2004; Uhlír et al., 1996).

C-type lectins are among molecules often involved in the immune system of arthropods (Ming et al., 2024). Arthropods generally lack a lymphocyte-mediated adaptive immune system, relying instead on their innate immunity, which consists of both cellular and humoral responses (Lavine and Strand, 2002; Lemaitre and Hoffmann, 2007). This system evolved over long-term co-evolution with pathogens (Sadd and Schimd-Hempel, 2006; Uvell and Engström, 2007). Some arthropod C-type lectins also exhibit functions beyond immunity, such as TcCTL1 from *Tribolium castaneum*, which is involved in embryogenesis (Zhang et al., 2023), or lectin Schlaff from *Drosophila melanogaster*, which influences the compactness of epidermal barrier (Zuber et al., 2019).

Recently, recombinant CTLs from several arthropod species have been successfully produced in bacterial expression systems. Examples include recombinant TgCTL-1 from the clam *Tegillarca granosa*, which plays a role in the recognition and binding of pathogens, and in the modulation of phagocytic activity of haemocytes (Ri et al., 2022); recombinant TcCTL3 from the insect *Tribolium castaneum,* which mediates the immune response via the pattern recognition, agglutination and antimicrobial peptides expression (Bi et al., 2019; Bi et al., 2020); and HlCLec from the Asian long-horned tick *Haemaphysalis longicornis* with three carbohydrate-binding domains, which participates in the tick’s defense against gram-negative bacteria (Maeda et al., 2016).

In our work, we focused on the production and characterization of one of the *I. ricinus* C-type lectins and its three domains to characterize binding properties and possibly uncover the function of this novel protein.

## Materials and methods

### Search for new lectins in the transcriptome of *Ixodes Ricinus*

Lectins of animal and bacterial origin with known structures or specificities and various functions were selected (Supplementary Table S1) and used to identify homologous proteins in the *Ixodes ricinus* transcriptome (GenBank: GIDG00000000). The search was carried out using the BioEdit program (Hall, 1999) which incorporates blastp (Altschul et al., 1990), and all six reading frames of the transcriptome were analysed.

Newly identified proteins were subjected to additional BLAST analyses and classified as potential lectins if a homologous lectin with at least 40% sequence identity and 60% sequence coverage was detected. Selected candidates were analysed further.

### Bioinformatic analysis of the putative lectin

To analyse the properties of the most promising predicted lectin, *I. ricinus* C-type lectin (IrCLec), we used ProtParam (Wilkins et al., 1999) for the prediction of physicochemical properties, WoLF PSORT (Horton et al., 2007) and DeepLoc 2.1 (Ødum et al., 2024) for the prediction of cellular localization, NetNGlyc (Gupta and Brunak, 2002) and NetOGlyc (Steentoft et al., 2013) for the detection of possible glycosylation sites, SignalP 6.0 (Teufel et al., 2022) for the determination of signal peptide presence, and DeepTMHMM (Hallgren et al., 2022) for the prediction of transmembrane regions. Sequence alignment was performed using EMBOSS Needle (Madeira et al., 2024). CDD database v.3.16 of NCBI (Marchler-Bauer et al., 2011), Pfam (El-Gebali et al., 2019), PROSITE (Sigrist et al., 2013), and ThreaDomEx (Wang et al., 2017) were used to search for conserved domains within the sequence. I-TASSER (Roy et al., 2010; Yang and Zhang, 2015; Zhang et al., 2017) was used for the prediction of binding sites.

Protein secondary and tertiary structures were predicted using AlphaFold 3 (Abramson et al., 2024). Structural models were generated for the full-length protein as well as for the putative carbohydrate recognition domains 1, 2, and 3 (CRD1, CRD2, and CRD3). The predicted structures were visualized using the PyMOL Molecular Graphics System (version 3.0.3; Schrödinger, LLC), and the presence and spatial localization of disulfide bridges were inferred from the three-dimensional models.

### Phylogenetic reconstruction

The full sequence of *I. ricinus* C-type lectin was used for the protein BLAST analysis to identify similar sequences among arthropod species (taxid:6656, accessed on 3 October 2024). Sequences sharing at least 30 % identity with IrCLec and covering at least 60 % of its length were selected for further analysis. Sequences with identity over 95 % percent were used only once and low quality and partial sequences were excluded. The phylogenetic tree was constructed using the Neighbour-Joining method (Saitou and Nei, 1987) based on MUSCLE alignment with MEGA 11 software, version 11.0.13 (Tamura et al., 2021). Bootstrap analysis was performed with 1000 replicates.

### Ticks

Adult females of *I. ricinus* were collected by flagging in a forest near Ceske Budejovice, Czech Republic, and kept at 95 % humidity, 24 °C, and 15/9 daylight settings. The adults were fed on guinea pigs by using plastic cylinders attached to guinea pigs’ backs. *I. ricinus* nymphs were supplied by the Animal facility, Biology Centre of Czech Academy of Sciences, Ceske Budejovice. The nymphs were fed on BALB/c mice (Charles River Laboratories) by using plastic cylinders attached to mice’s backs. All experiments were conducted in accordance with the animal protection law of the Czech Republic (§17, Act No. 246/1992 Sb) with the approval of Czech Academy of Sciences (approval no. 79/2013).

### qRT-PCR tissue and stage profiling of IrCLec

For the tissue profiling, unfed, semi-engorged (five days of feeding), and fully fed females (four days after engorgement) were dissected under the stereo microscope for various tissues as described previously (Honig Mondekova et al., 2017). For the stage profiling, unfed and fed larval, nymphal, and adult ticks, as well as eggs and unfed males, were homogenized. The material used for gene expression analysis 12 hours after pathogen injection (*Escherichia coli*, *Micrococcus luteus*, *Candida albicans*, *Borrelia afzelii*) in unfed adults was obtained as described previously (Urbanová et al., 2015). All samples were prepared in biological triplicates. Total RNA was extracted using NucleoSpin RNA II kit (Macherey-Nagel) and its integrity was assessed by agarose gel electrophoresis. The RNA was reverse transcribed (0.5 μg per reaction) into cDNA using the Transcriptor High-Fidelity cDNA Synthesis Kit (Roche) and diluted 20-fold in sterile water. Gene expression was determined by quantitative real-time PCR (qRT-PCR) using a LightCycler 480 (Roche) and SYBR green chemistry using primers listed in Table 1. Relative expression was normalized to *I. ricinus elongation factor 1* (GU074769) according to the mathematical model of Pfaffl (Pfaffl, 2001).

**Table 1.**
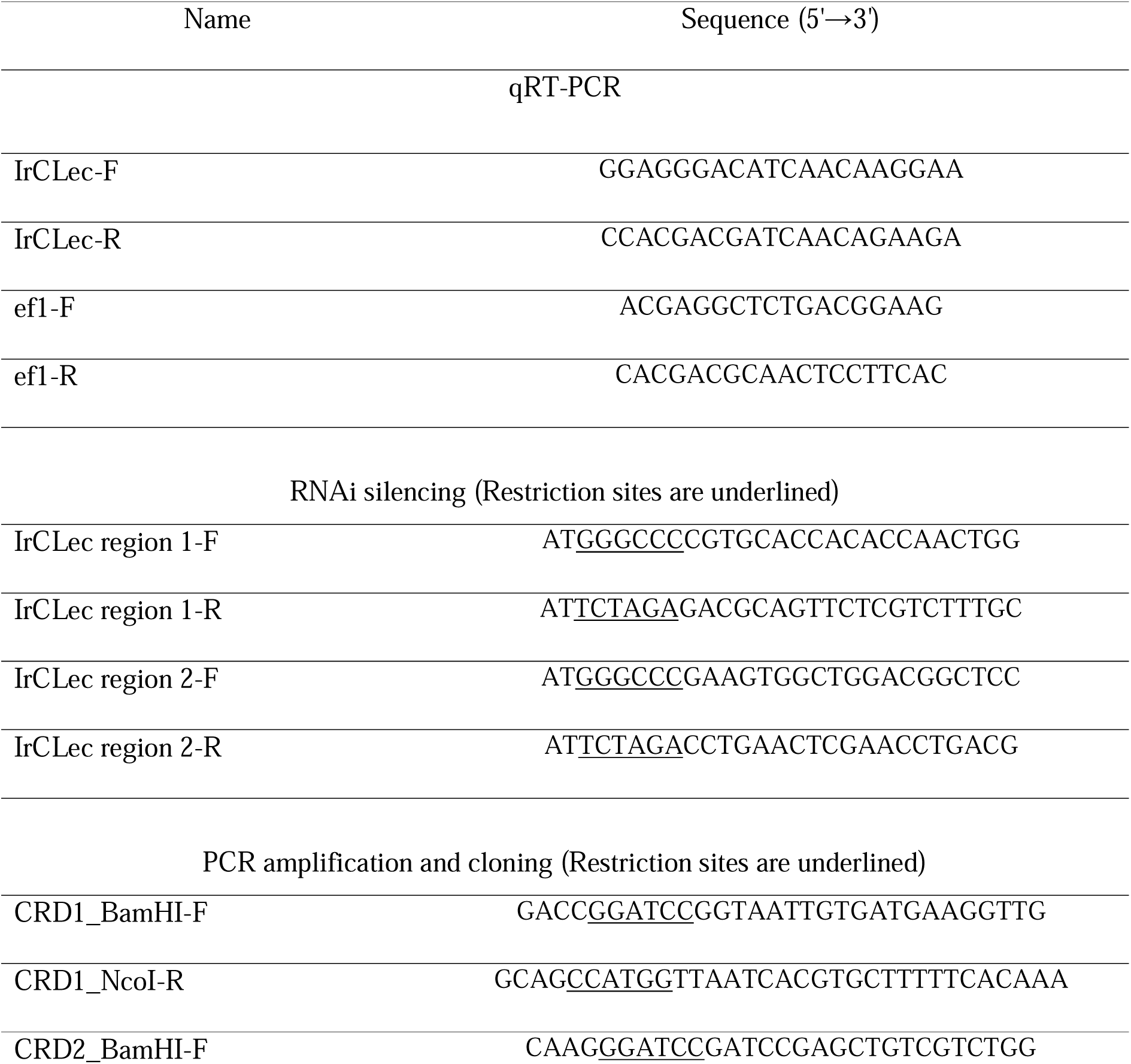
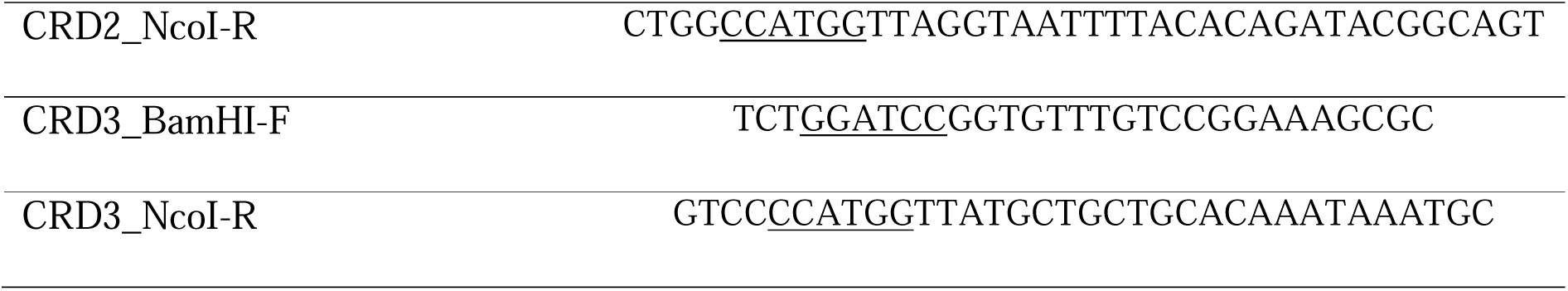
List of used primers.

### RNA silencing

Two fragments of the *IrCLec* gene, position 428-809 (IrCLec region 1) and 880-1367 (IrCLec region 2) of GIDG01027020, were cloned into the pll10 vector using primers listed in Table 1 and the dsRNA synthesized as described before (Hajdušek et al., 2009). The dsRNA (3 µg/µL) was injected into the hemocoel of nymphs (32.2 nL) using Nanoinject II (Drummond) injector. After three days of rest, the injected ticks were fed on BALB/c mice. The level of gene silencing was checked by qRT-PCR in a mix of five fully fed nymphs. The gene silencing data were analysed using GraphPad Prism 4.0 (GraphPad Software) employing the Mann-Whitney test. All results are expressed as the mean ± standard error (SEM). Gene expression profiling data (qRT-PCR) represent three biological replicates and were not analysed by statistical methods.

### Production of recombinant domains of IrCLec

The coding part of *IrCLec* gene, codon-optimized for *E. coli* production, was obtained in pMT plasmid from GeneArt Gene Synthesis (Thermo Fisher Scientific). Sequences of each predicted carbohydrate recognition domain (CRD1, CRD2, and CRD3) were amplified using PfuUltra High-Fidelity DNA polymerase (Agilent Technologies) and specific primers (Table 1) and ligated into the expression vector pRSET-A (Invitrogen) using BamHI and NcoI restriction sites.

The newly prepared vectors were transformed into *Escherichia coli* strain Rosetta-gami 2(DE3)pLysS (Novagen) for protein expression. Protein production was induced by isopropyl-beta-D-thiogalactopyranoside (IPTG) at a final concentration of 1 mM. The CRD3 domain was produced for 3 hours at 37 °C and CRD1 and CRD2 were produced for 18 hours at 18 °C.

Cells from 1 l of LB medium were lysed by sonication in buffer A (20 mM Tris base, pH 7.5, 150 mM NaCl and 0.5 mM CaCl_2_). The recombinant proteins were purified from the lysate by affinity chromatography using the Ni-NTA Superflow Cartridge 5 mL (Qiagen) with buffer A. Buffer B (20 mM Tris base, pH 7.5, 150 mM NaCl, 0.5 mM CaCl_2_, and 0.5 M imidazole) was used for gradient elution under non-denaturing conditions.

The purified proteins were dialysed back to buffer A and concentrated using a Vivaspin 20 centrifugal concentrator (Sartorius Stedim Biotech). Protein concentration was measured by NanoDrop 1000 Spectrophotometer (Thermo Fisher Scientific) at the wavelength 280 nm using the molar extinction coefficient of analysed proteins. The obtained proteins were detected by reducing 15 % sodium dodecyl sulphate-polyacrylamide gel electrophoresis (SDS-PAGE) at 120 V for 90 min and visualized with Coomassie Brilliant Blue R-250.

### Western blotting analysis

Proteins were transferred from the 15 % polyacrylamide gel to a nitrocellulose membrane (0.2 μm, Bio-Rad). After transfer, the membrane was blocked for 1 hour at room temperature using 5 % skim milk in Tris-buffered saline (TBS) (20 mM Tris base, 150 mM NaCl, pH 7.6). Then, the membrane was washed three times with TBS + 0.05 % Tween20 (TBST) and incubated with Monoclonal Anti-polyHistidine-Peroxidase antibody produced in mouse (Sigma-Aldrich) at a 1:2000 dilution in TBST with 1 % bovine serum albumin overnight at 4 °C. After incubation, membranes were washed twice with TBST and once with TBS. The detection was performed using a horse radish peroxide substrate solution (15 mL of TBS, 3 mL of methanol, 3 mg of 4-chloro-1-naphthol, and 50 µL of 30 % hydrogen peroxide solution). The protein markers used were PageRuler™ Plus Prestained Protein Ladder (Thermo Fisher Scientific) and Protein Marker III 6,5-200 kDa (AppliChem).

### Hemagglutination assay and hemagglutination inhibition assay

Hemagglutination and hemagglutination inhibition assays were performed using anonymized human erythrocytes from healthy donors of blood groups A, B and O (purchased from the Department of Transfusion & Tissue Medicine, University Hospital Brno, Czech Republic). The red blood cells (RBCs) were washed three times with buffer A, centrifuged at 2000 g for 3.5 minutes and diluted to a 50 % solution by the same buffer. RBCs were stabilized by 0.01 % (w/v) NaN_3_ and stored at 4 °C. Hemagglutination experiments were performed in a 96-well U-shaped microtiter plate (Anicrin). Each well (except the first well in the row) was filled with 50 μL of buffer A. 100 μL of protein sample was pipetted into the first well and serially diluted by two-fold dilution series, except for the last well (negative control). 50 μL of the 2 % RBCs suspension was added to each well and mixed. The mixture was incubated for 1 hour at room temperature, and the formation of aggregates was analysed. The highest dilution of the protein solution showing complete agglutination (the titter) was determined visually.

For the hemagglutination inhibition assay, 25 μL of the recombinant His-CRD1 or His-CRD2 was mixed with 25 μL of serially diluted saccharides (α-L-fucose (L-Fuc), α-D-galactose (D-Gal), *N*-acetyl-α-D-galactosamine (GalNAc) (Carbosynth Limited), α-L-galactose (L-Gal), α-D-glucose (D-Glc), α-D-mannose (D-Man) (Duchefa Biochemie) dissolved in buffer A. Mixtures were incubated for 15 min at room temperature. The final concentrations of L-Fuc, D-Gal, L-Gal, D-Glc, D-Man, and GalNAc ranged from 12.5 mM to 0.024 mM. The final concentration of the recombinant His-CRD1 or His-CRD2 was the lowest concentration causing full hemagglutination times 2. Subsequently, 50 μL of the 2 % RBC suspension was added to each well, mixed, and incubated for 1 hour. The minimal concentration of saccharides showing complete inhibition of hemagglutination (minimal inhibitory concentration, MIC) was determined visually.

### Glycan array

His-tag purified proteins CRD1, CRD2, and CRD3 were labelled with Monolith NT-His-Tag labelling Kit (RED-tris-NTA 2^nd^ generation dye, NanoTemper Technologies) in a 1:1 molar ratio (the dye was prepared according to the manual). The glycan microarray chip containing 381 mammalian and bacterial glycans in hexaplicates (Semiotik; chip format OS090418) was incubated in 2.0 mL of buffer A supplemented with 0.1 % (v/v) Tween 20 and placed in a humid chamber for 15 min (25 °C, 35 rpm). Then, the buffer was replaced by 0.5 mL of labelled studied protein (0.2 μM) diluted in buffer A with 0.1 % Tween 20 and 1 % (w/v) bovine serum albumin (Serva). After incubation (60 min, 25 °C, 35 rpm), the chip was washed with decreasing concentrations (100 % and 50 %) of buffer A with 0.1 % Tween 20, followed by multiple immersions in ultrapure water. Subsequently, the chip was dried by air blow. Fluorescence was read with a scanner InnoScan 1100 AL (Innopsys) with a 635 nm laser at 20 °C. The data were analysed using Mapix 8.2.2 software and an online glycan chip converter (Semiotik, https://rakitko.shinyapps.io/semiotik). Ligand is considered significant when a minimum of five spots from the hexaplicate have SNR higher than 3, and the median signal obtained for the ligand is at least three times higher than the negative control defined by the manufacturer. The raw glycan-array data are presented in Supplementary Tables S2, S3, S4.

## Results

### Identification and characterization of IrCLec

During a search for novel lectins in the previously sequenced *Ixodes ricinus* transcriptome (Vechtova et al., 2020), a novel C-type lectin, designated IrCLec, was identified. IrCLec showed sequence homology to lung surfactant protein from *Rattus norvegicus*, with 26% sequence identity and 46% similarity. Subsequent bioinformatic analyses identified a closer homolog, HlCLec, a C-type lectin from the Asian tick *Haemaphysalis longicornis*, with 95% sequence coverage and 49% sequence identity. The amino acid sequence of IrCLec is 519 amino acids long, with a calculated molecular weight of 57.5 kDa and an isoelectric point of 6.02. Bioinformatic analysis of the sequence revealed the presence of three carbohydrate recognition domains (CRD1, CRD2, and CRD3) as well as an N-terminal signal peptide and a C-terminal transmembrane helix, matching the arrangement of HlCLec (Figure 1). The domains have a variable level of similarity to each other, the similarity of CRD1 to CRD2 being 38.9 %, CRD1 to CRD3 49.6 % and CRD2 to CRD3 30.5 %. Potential carbohydrate-binding and calcium-binding motifs were identified for all three domains. The sequence of CRD3 contains the typical binding motifs of C-type lectins – EPN and WND. The other two domains appear to contain non-common binding motifs: QPN and WNQ in the CRD1, and QPG and WSV in the CRD2. WoLF PSORT and DeepLoc predicted that IrCLec is a membrane protein. Two potential *N*-glycosylation sites (N43, N346) and three potential *O*-glycosylation sites (T252, S296, S446) were detected.

**Figure 1.**
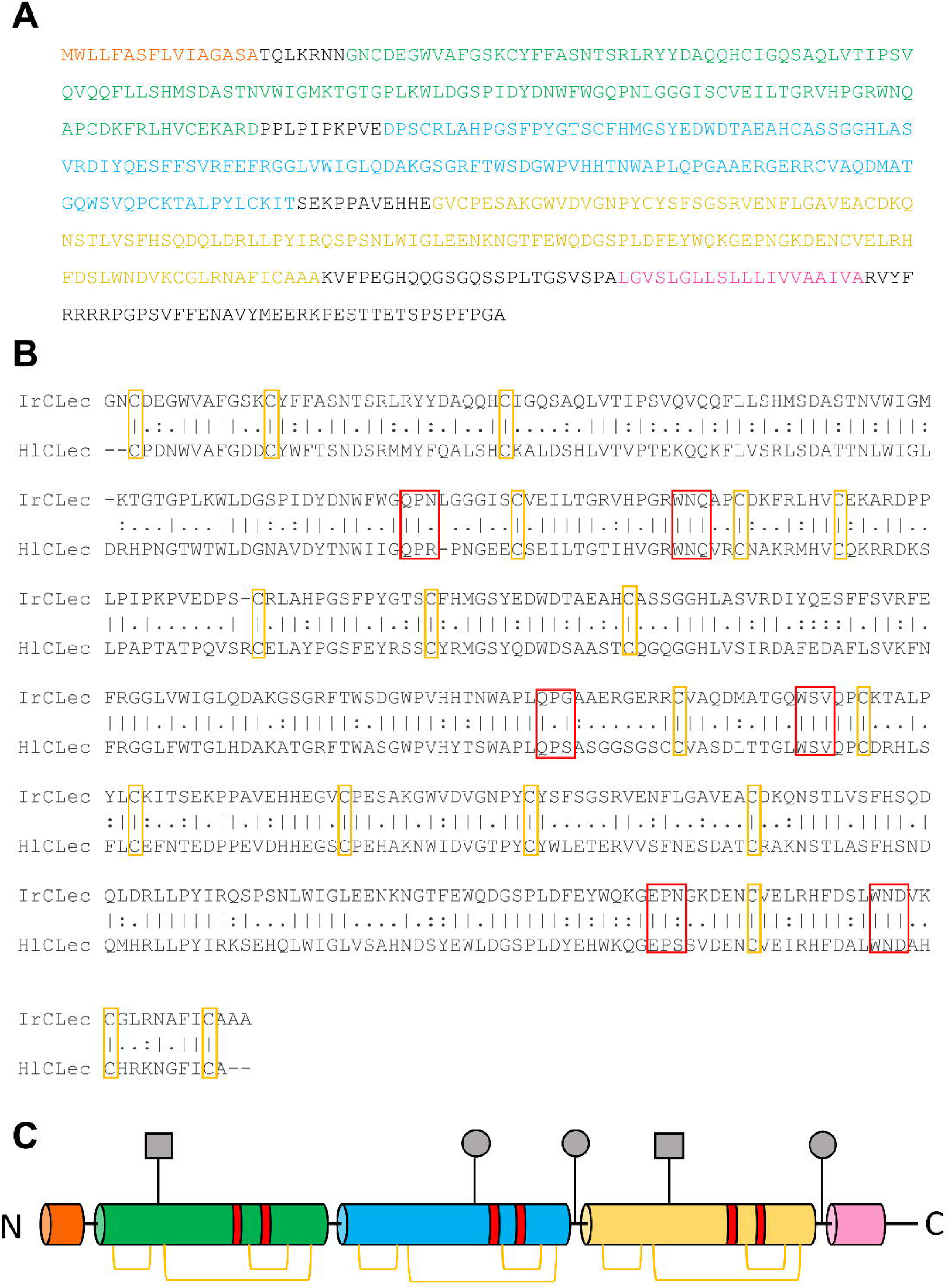
The sequence of IrCLec. **A** The signal peptide is coloured orange, carbohydrate recognition domain 1 (CRD1) is coloured green, CRD2 is coloured blue, CRD3 is coloured yellow, and the transmembrane region is coloured pink. **B** Alignment of the IrCLec and HlCLec proteins from the beginning of first domain to the end of the third domain. Orange frames mark cysteines, red frames mark carbohydrate-binding motifs. **C** Scheme of IrCLec structure. The signal peptide is coloured orange, CRD1 is coloured green, CRD2 is coloured blue, CRD3 is coloured yellow, the transmembrane region is coloured pink, carbohydrate-binding motifs are coloured red. N indicates N-terminus, C indicates C-terminus, orange brackets indicate disulfide bridges, grey squares indicate N-linked glycosylation, and grey circles indicate O-linked glycosylation.

### The prediction of tertiary structure

The tertiary structures of individual IrCLec domains and the full-length protein were predicted using AlphaFold 3 (Figure 2). The predicted pTM scores were 0.91 for CRD1, 0.88 for CRD2, 0.88 for CRD3, and 0.68 for the full-length IrCLec protein. All three CRDs were predicted to adopt the characteristic fold of the C-type lectin-like domain superfamily, comprising a double-loop (“loop-in-a-loop”) structure stabilized by conserved disulfide bridges located at the bases of the loops (Zelensky and Gready, 2005). According to I-TASSER predictions, the extended loop regions of the CRDs may be involved in carbohydrate binding. These regions spatially correspond to the potential carbohydrate-binding motifs QPN and WNQ in CRD1, QPG and WSV in CRD2, and EPN and WND in CRD3.

**Figure 2.**
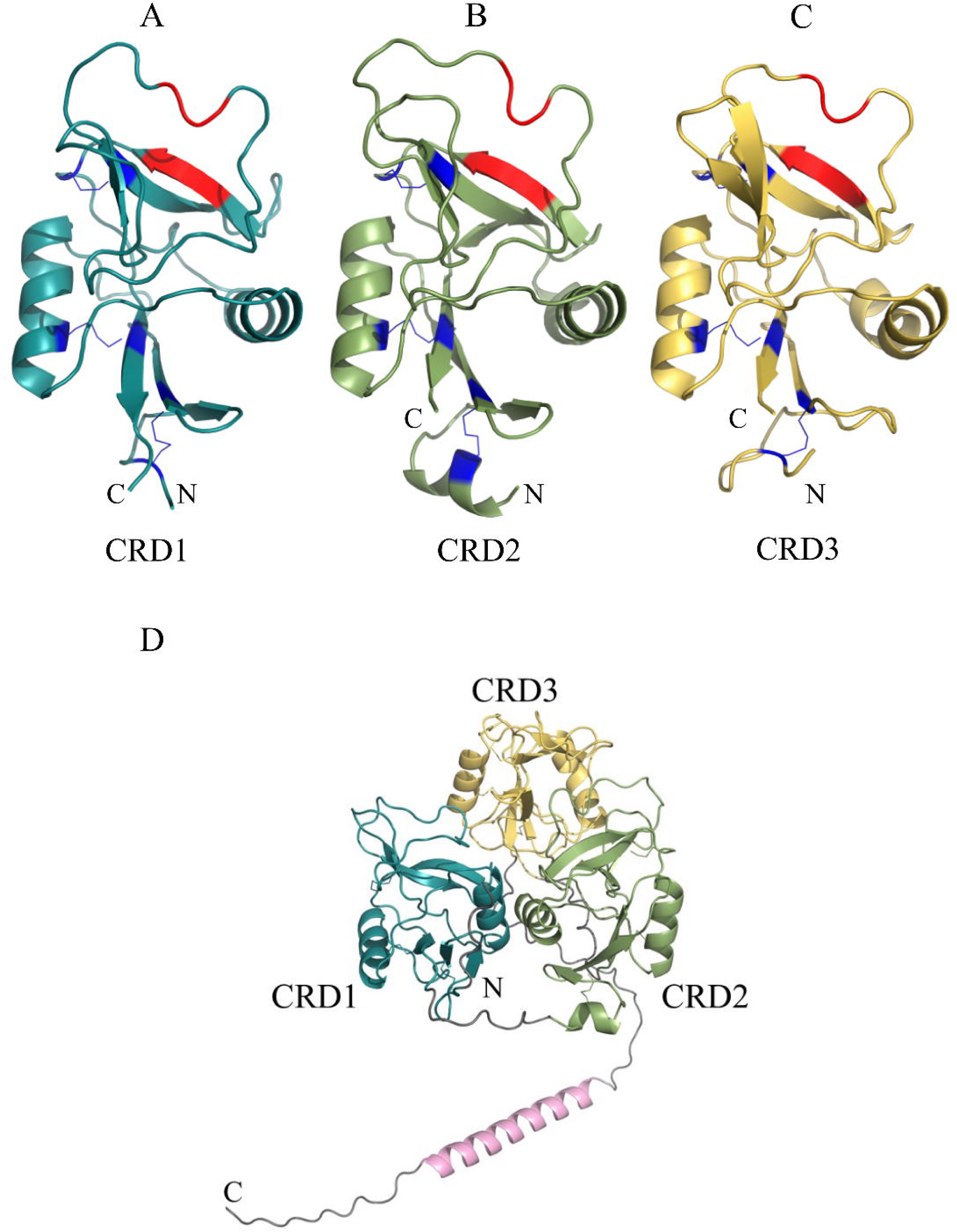
The 3D structures of putative CRD1 (**A**), CRD2 (**B**), and CRD3 (**C**) domains predicted by AlphaFold 3: potential carbohydrate-binding motifs are coloured red, and disulfide bridges are coloured orange. **D** Predicted structure of the whole IrCLec.

### Phylogenetic reconstruction

Sequence homologs of IrCLec were identified across several arthropod species. With the exception of HlCLec, none of the identified homologs have been functionally or structurally characterized to date. The closest homolog of IrCLec was identified as a putative macrophage mannose receptor 1 from *Ixodes scapularis* (Figure 3). Based on annotation, more than half of the homologous sequences were classified as macrophage mannose receptors, four were annotated as hypothetical proteins, three as secretory phospholipase A2 receptors, and one as a brevican core protein. All annotated protein types contain at least one C-type lectin domain within their predicted domain architecture. Phylogenetic analysis revealed that homologs from tick species form a distinct clade, representing the phylogenetically closest group to IrCLec. The remaining homologs originate from a range of arthropod taxa, including spiders, crabs, prawns, as well as single representatives from lobster, amphipod, and scorpion species.

**Figure 3.**
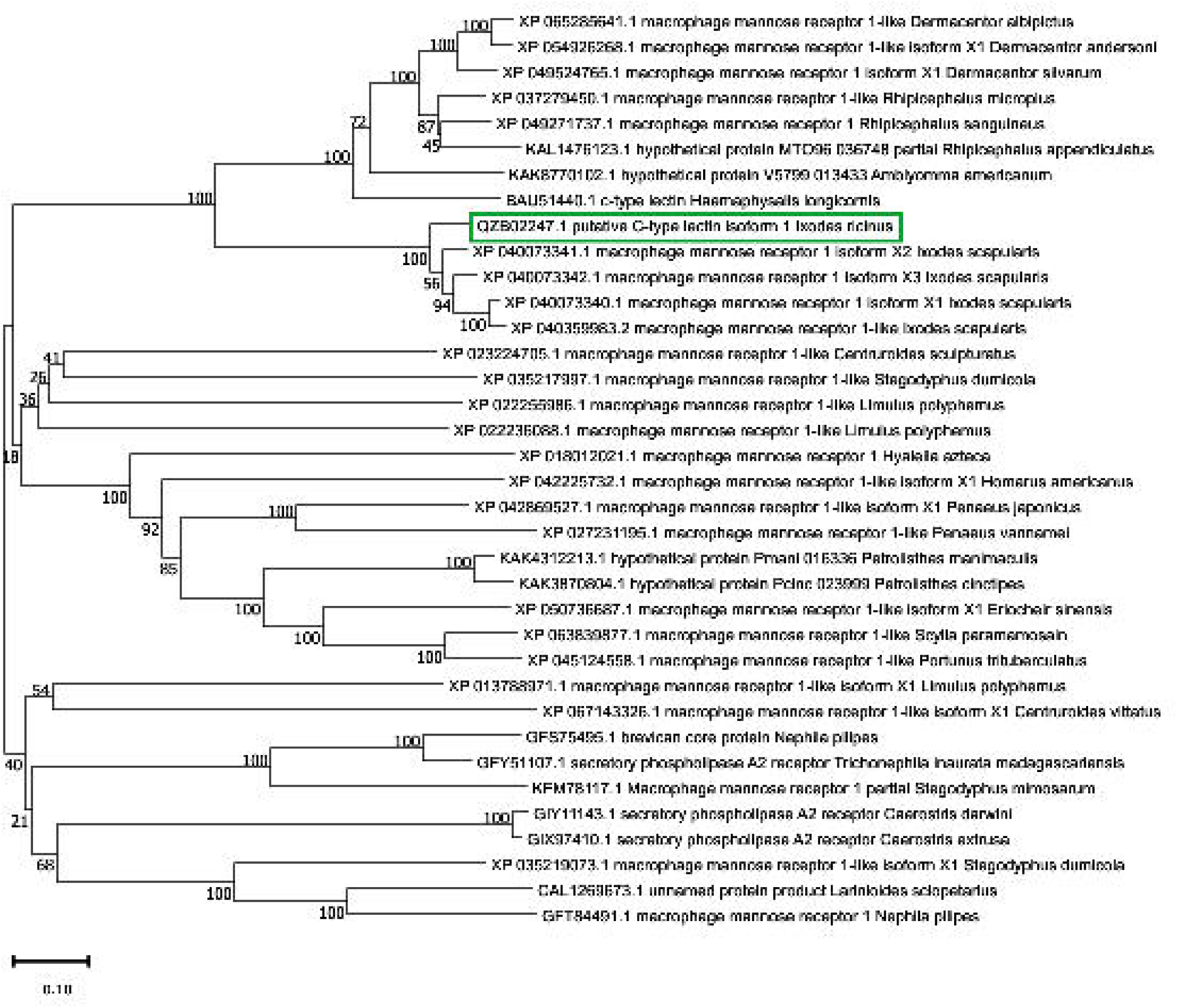
Phylogenetic tree showing the relationship between IrCLec and related arthropod amino acid sequences, reconstructed using the Neighbour-Joining method and based on alignment using MUSCLE. Bootstrap analysis was performed with 1000 replicates, and the numbers at the nodes represent bootstrap values. IrCLec is highlighted by a green frame.

### Determination of biological functions of IrCLec in ticks

To assess IrCLec the biological functions of IrCLec in tick, tissue- and stage-gene expression profiling was performed using qRT-PCR, followed by functional analysis using RNA interference in tick nymphs. IrCLec expression was detected predominantly in the midgut before, during, and after blood feeding (Figure 4A). Lower but detectable expression levels were also observed in haemocytes and ovaries. In addition, IrCLec expression was stimulated by blood feeding in tick larvae and nymphs.

**Figure 4.**
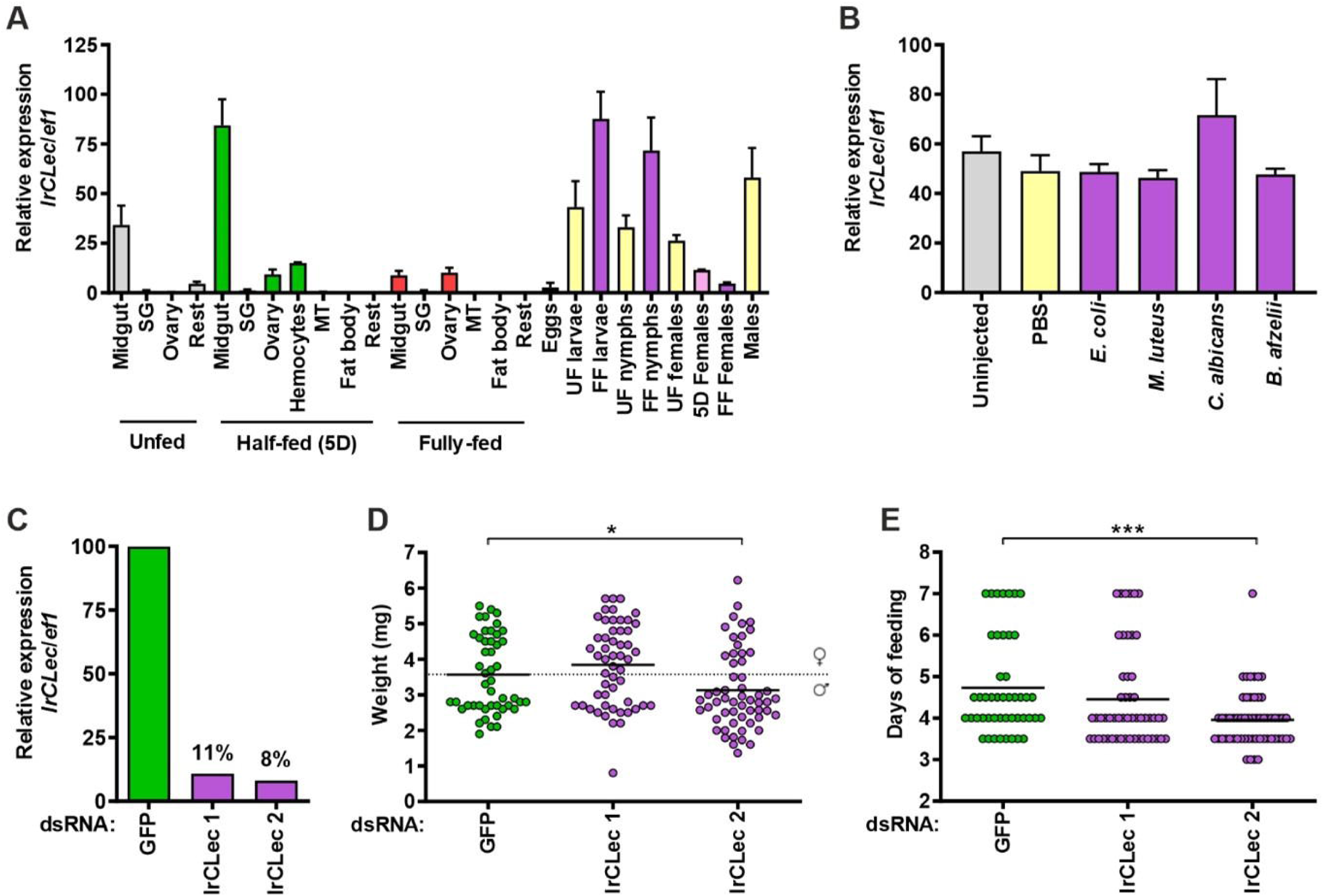
Biological functions of IrCLec in *I. ricinus*. **A** Relative expression of *IrCLec* in the tick tissues of unfed, half-fed (five days of feeding), and fully-fed (four days after engorgement) adult females and expression in the various tick stages. SG - salivary glands, MT - Malpighian tubules, UF - unfed, FF - fully-fed, 5D - five days, *ef1* - *elongation factor 1* (housekeeping gene). The samples were prepared in biological triplicates. In each graph, cDNA with the highest expression was set as 100 %. **B** Relative expression of *IrCLec* in unfed adult females 12 hours after injection of pathogens. **C** Relative expression of *IrCLec* in the dsRNA-treated fully fed nymphs (control of gene silencing). Two dsRNAs covering diverse gene regions were prepared for *IrCLec* (denoted as 1 and 2). gfp dsRNA was used as a control. D Effect of gene silencing on the weight of fully fed nymphs. The dotted line depicts the theoretical border between females (heavier) and males. The values in each group represent a mixture of ticks fed on three mice (20 nymphs were fed on individual mice). E Effect of gene silencing on the duration of nymph feeding. *P ≤ 0.05; ***P ≤ 0.001.

Because tick lectins have been previously described as potential immune molecules (Honig Mondekova et al., 2017), *IrCLec* gene expression was measured 12 h after injection of several pathogens. No significant changes in *IrCLec* expression were observed following either PBS injection (effect of injury) or pathogen challenge (Figure 4B).

Next, gene silencing of *IrCLec* was studied using RNA interference in tick nymphs. Two independent dsRNAs targeting different regions of the *IrCLec* gene were synthesized and injected into nymphs. The expression of *IrCLec* decreased by 89% and 92% for dsRNA targeting regions 1 and 2, respectively (Figure 4C). After dsRNA injection, nymphs were allowed to feed on BALB/c mice. Silencing of *IrCLec* using dsRNA targeting region 2 significantly diminished engorgement weights of nymphs and reduced feeding time (Figures 4D, 4E). No significant differences in moulting success were observed between control and IrCLec-silenced nymphs, with moulting rates of 60%, 71%, and 60% for dsGFP, dsIrCLec region 1, and dsIrCLec region 2, respectively. The engorged nymphs did not differ in the moulting efficacy compared with the dsGFP control nymphs (60 %, 71 %, and 60 % of nymphs moulted in *dsGFP*, *dsIrCLec* 1 and *IrCLec* 2, respectively).

### Production of carbohydrate recognition domains of IrCLec

The individual carbohydrate recognition domains of IrCLec were separated at the gene level and cloned individually into the pRSET A expression vector. Recombinant His-tagged domains were produced in *Escherichia coli* Rosetta-gami 2(DE3) pLysS cells and purified using Ni–NTA affinity chromatography. Production and purification of all three domains were confirmed by SDS–PAGE and Western blot analysis (Figure 5). Western blot analysis revealed basal production of all recombinant proteins in uninduced cells. All three domains were produced predominantly in the insoluble form, with only a minor proportion detected in the soluble fraction. Despite the low solubility, purification of soluble protein fractions was successful, yielding highly purified recombinant proteins. The yield of soluble protein was approximately 0.1 mg per litre of bacterial culture for each domain.

**Figure 5.**
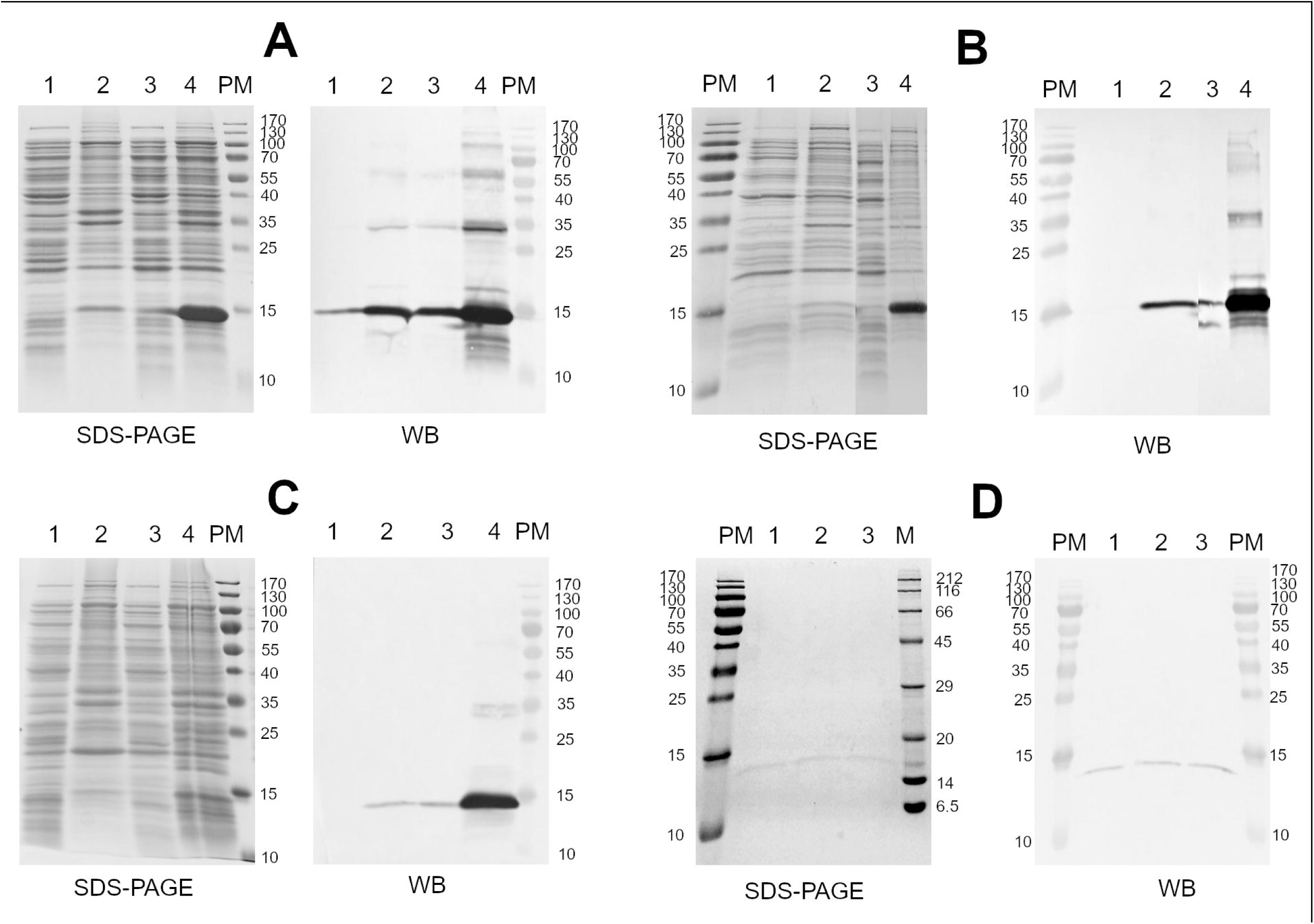
The production of recombinant CRDs in bacterial cells. Cell lysates containing His-CRD1 (**A**), His-CRD2 (**B**), and His-CRD3 (**C**) were separated by the reducing 15 % sodium dodecyl sulphate-polyacrylamide gel electrophoresis (SDS-PAGE) and visualized with Coomassie Brilliant Blue R-250. Monoclonal Anti-polyHistidine-Peroxidase antibodies were used to detect the His-tagged proteins on the nitrocellulose membranes by Western blotting (WB). 1 – cell lysates before induction, soluble fraction; 2 – cell lysates before induction, insoluble fraction; 3 – cell lysates after 1 mM IPTG induction, soluble fraction; 4 – cell lysates after 1 mM IPTG, insoluble fraction; PM – prestained protein marker. The purity of Ni-NTA purified proteins (**D**) was confirmed by SDS-PAGE and WB. M – protein marker, 1 - His-CRD1; 2 - His-CRD2; 3 - His-CRD3.

### Hemagglutination activity of separate His-tagged CRDs of IrCLec

Hemagglutination activity of individual IrCLec carbohydrate recognition domains was tested using human red blood cells (RBCs) of blood groups A, B, and O (Figure 6). The recombinant protein His-CRD3 did not exhibit hemagglutination activity at the tested concentrations (Table 2). In contrast, His-CRD1 and His-CRD2 were able to hemagglutinate all three blood groups. His-CRD1 exhibited the strongest hemagglutination activity toward blood group B, followed by blood groups A and O. His-CRD2 showed the highest preference for blood group O, followed by blood groups A and B. No hemolysis of RBCs was observed under any of the tested conditions.

**Figure 6.**
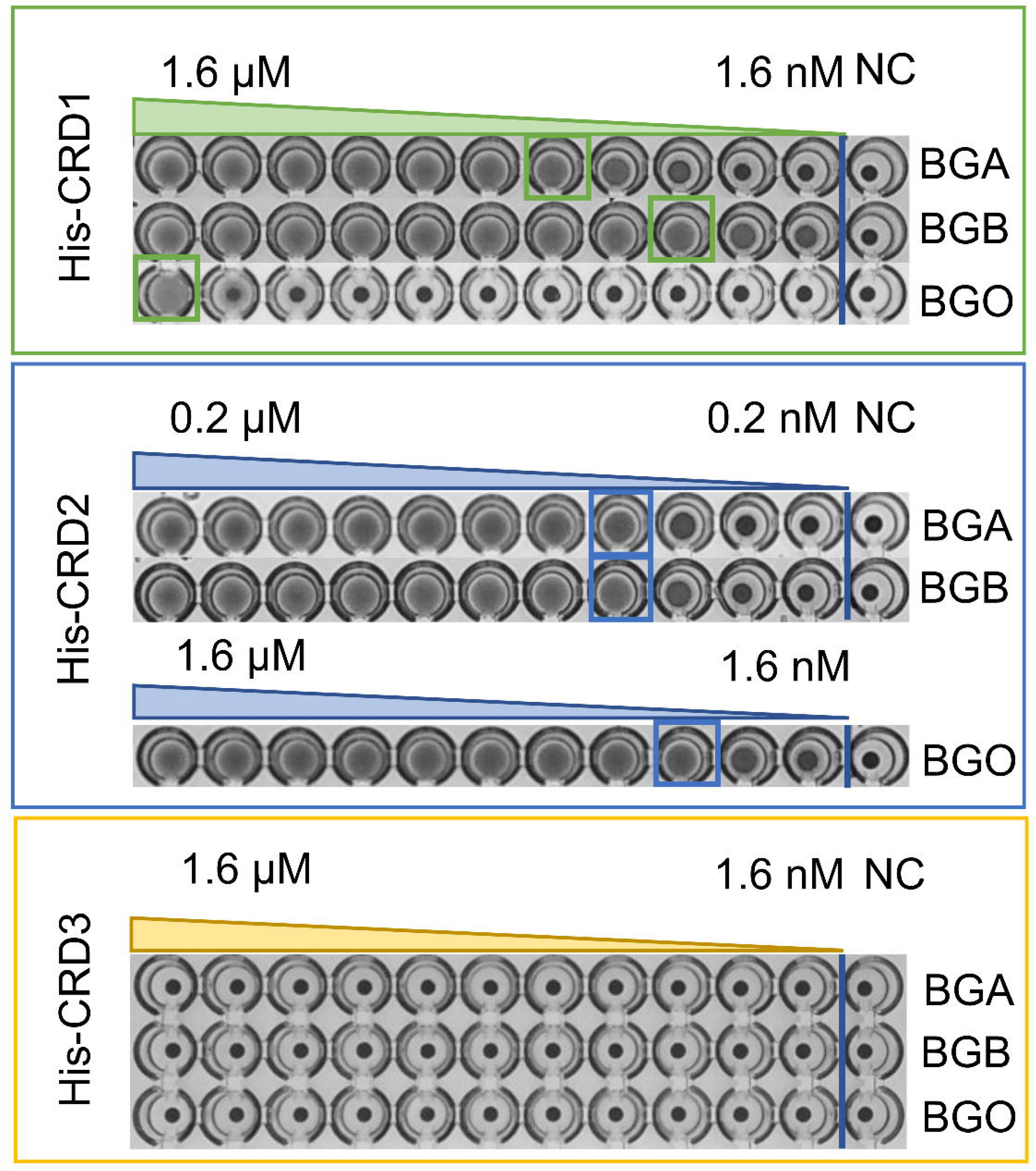
Hemagglutination assay of His-tagged CRDs. Protein concentration decreases two-fold from left to right. Agglutinated red blood cells form a diffuse mat, whereas non-agglutinated red blood cells sediment and form a clear dot at the bottom of the well. The wells marked as NC (negative control) represent control experiments in the absence of proteins. Titter is marked by a frame. BGA – blood group A, BGB – blood group B, BGO – blood group O.

**Table 2.** Hemagglutination activity of His-tagged CRD1, CRD2, and CRD3.

| Protein | BGA | BGB | BGO |
| --- | --- | --- | --- |
|  | Titre of HA, nM |  |  |
| His-CRD1 | 25 | 6.25 | 1600 |
| His-CRD2 | 16 | 16 | 6.25 |
| His-CRD3 | ND | ND | ND |

ND - not detected

### Hemagglutination inhibition assay

Preliminary carbohydrate-binding specificity of His-CRD1 and His-CRD2 was examined using a hemagglutination inhibition assay. The monosaccharides D-Man, D-Glc, D-Gal, and D-GalNAc did not inhibit hemagglutination (Table 3). In contrast, monosaccharides L-Fuc and L-Gal exhibited inhibitory effects.

**Table 3.** Minimal inhibitory concentration (MIC) values for inhibition of hemagglutination caused by His-CRD1 and His-CRD2, potency (MIC of L-Fuc/MIC of sugar), and β-value (potency/valency) of tested sugars with blood groups A and B.

| Inhibitor | MIC<br>(mM) | Potency | Valency | β | MIC<br>(mM) | Potency | Valency | β |
| --- | --- | --- | --- | --- | --- | --- | --- | --- |
|  | BGA |  |  |  | BGB |  |  |  |
| CRD1 |  |  |  |  |  |  |  |  |
| L-Fuc | 6.25 | 1 | 1 | 1 | ND | - | 1 | - |
| L-Gal | 12.5 | 0.5 | 1 | 0.5 | ND | - | 1 | - |
| D-Man, D-Glc,<br>D-Gal, GalNAc | ND | - | 1 | - | ND | - | 1 | - |
| CRD2 |  |  |  |  |  |  |  |  |
| Fuc | 12.5 | 1 | 1 | 1 | ND | - | 1 | - |
| L-Gal, D-Man,<br>D-Glc, D-Gal,<br>GalNAc | ND | - | 1 | - | ND | - | 1 | - |

ND - not determined

For blood group B, no inhibitory effect of monosaccharides was observed for either domain. For blood group A, both L-Fuc and L-Gal were able to inhibit the hemagglutination of His-CRD1, with L-Fuc having a minimal inhibitory concentration two times lower than L-Gal. For hemagglutination of blood group A by His-CRD2, inhibition was observed exclusively with L-Fuc.

### Glycan array

Binding specificities of His-CRD1, His-CRD2, and His-CRD3 were further analysed using a glycan array microchip containing 381 immobilized ligands representing various bacterial and mammalian glycans (Figure 7).

**Figure 7.**
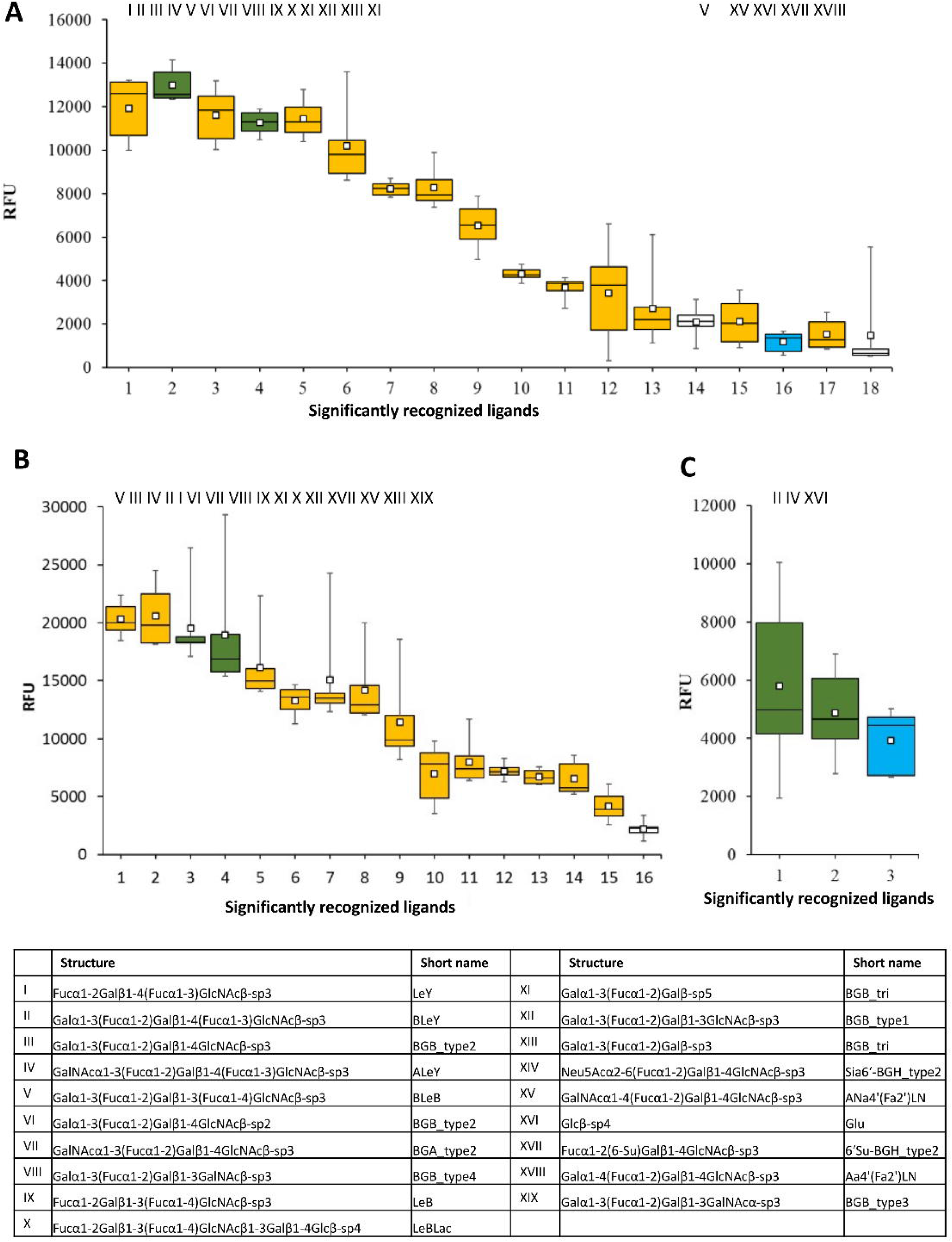
Box and whisker plot for glycan array screening with His-CRD1 **(A)**, His-CRD2 **(B)**, and His-CRD3 **(C)** domains labelled with Monolith His-Tag Labelling Kit (RED-tris-NTA 2^nd^ generation dye). The Y-scale represents the relative fluorescence units (RFU). The bottom and top of the box represent the first and third quartiles, the band inside the box is the median, the small square inside the box represents the mean, and the ends of the whiskers represent the minimum and maximum values of the obtained data. Glycans recognized by all three domains are coloured green, glycans recognized by His-CRD1 and His-CRD2 are coloured orange, glycans recognized by His-CRD1 and His-CRD3 are coloured blue, glycans recognized only by one domain are coloured white. The structures and the short names of ligands are given in the table. The linkers are as follows: sp2, −O(CH_2_)_2_NH_2_; sp3, −O(CH_2_)_3_NH_2_; sp4, −NHCOCH_2_NH_2_; sp5, −O(CH_2_)_3_NH-CO(CH_2_)_5_NH_2_.

With the one exception of β-D-glucose, all recognized ligands corresponded to derivatives of human blood group antigens, including blood groups A, B, and O, as well as Lewis b and Lewis Y antigens. These glycans share a common structural motif, Fucα1–2Gal.

The blood group–related pentasaccharides ALeY (GalNAcα1–3(Fucα1–2)Galβ1–4(Fucα1–3)GlcNAcβ) and BLeY (Galα1–3(Fucα1–2)Galβ1–4(Fucα1–3)GlcNAcβ) were recognized by all three domains. His-CRD1 recognized in total 18 ligands, whereas His-CRD2 bound to 16 ligands, of which 15 were shared with His-CRD1. In contrast, His-CRD3 showed significant binding to only three ligands: ALeY, BLeY, and β-D-glucose.

## Discussion

Lectins are specific carbohydrate-binding proteins involved in various recognition processes. Lectins from pathogens often function as virulence factors mediating adhesion to host cells and contributing to infection development. However, lectins also play roles in the immune system, establishment of symbiosis, growth regulation and differentiation, they serve as storage proteins or mediate glycoprotein folding.

C-type lectin family is large, diverse and widespread among animal species. Many CTL-containing receptors are integral components of the immune system. In this study, we describe a novel C-type lectin from the tick *Ixodes Ricinus* discovered in its transcriptome (IrCLec).

IrCLec displays sequence similarity to macrophage mannose receptors (MRs) from several tick species as well as other arthropods; the closest homologues being the macrophage mannose receptor isoforms from *Ixodes scapularis*. MRs are also expressed in mammals by macrophages, dendritic cells, and specific lymphatic or endothelial cells, and play an important role in immune homeostasis (Azad et al., 2014). In humans, this protein mediates endocytosis of glycoproteins by macrophages and acts as phagocytic receptor for bacteria and fungi. It is also involved in HIV binding and transmission as well as recognition of other viruses, including dengue virus, influenza virus and hepatitis B virus (Brown et al., 2018; Monteiro and Lepenies, 2017; Op den Brouw et al., 2009;). Notably, all MRs sequence homologues of IrCLec are currently annotated as predicted or hypothetical proteins. However, the only characterized protein among IrCLec homologues, *Haemaphysalis longicorni*s C-lectin (HlCLec), was also suggested to play a role in the immune defense, particularly against Gram-negative bacteria. Therefore, we initially hypothesized that the C-type lectin from *I. ricinus* may also contribute to immune defense, possibly as a recognition protein for carbohydrates on the surface of pathogens. This was also supported by localization prediction results which classified IrCLec as an integral membrane protein.

However, injections of pathogens into ticks did not have any observable effect on the expression level of the *IrCLec* gene, barring a slight expression increase in response to the presence of *C. albicans*. These results do not completely disprove the hypothesis of immune functions of IrCLec, as it belongs to surface receptors and the levels of its expression might simply not directly respond to bacterial infection. Generally, the expression of IrCLec was observed predominantly in the tick’s midgut with expression levels significantly increasing after blood-feeding. RNAi silencing of the *IrCLec* gene significantly diminished engorgement weights of *I. ricinus* nymphs and reduced feeding time, while moulting efficacy of engorged nymphs was not affected. These findings indicate that another, or possibly even primary, role of the protein could be the facilitation or regulation of blood intake and digestion in the midgut.

C-type lectins of invertebrate having a role in the feeding process of invertebrates is unusual, as most reports indicate functions in the immune system (Ming et al., 2024, Xia et al., 2018). However, few exceptions have been described. In the oyster *Crassostrea virginica*, lectins CvML3912 and CvML3914 have been found to play a role in efficient food particle sorting. These lectins were confirmed to be present in feeding organs, and *C. virginica* is able to distinguish nutritious particles from other matter through their interaction with saccharides of microalgae (Espinosa and Allam, 2018). A similar role was described for the C-type lectin Rpcl in *Ruditapes philippinarum* (Chen et al., 2023). It is therefore possible, that the function of IrCLec is also an interaction with food, in this case blood cells and the lectins function indeed differs from the close homologue HlCLec.

For further functional analysis, interactions of IrCLec with red blood cells were studied. Hence all three CRDs were produced in recombinant form as separate His-tagged proteins. The production and purification were successful, albeit the yields were modest. No hemolytic activity was detected, indicating that IrCLec does not function as a hemolysin. Notably, His-CRD1 and His-CRD2 exhibited hemagglutination activity, whereas His-CRD3 did not. Both His-CRD1 and His-CRD2 showed stronger hemagglutination toward blood groups A and B compared to blood group O, and their activity against BGA was inhibited by L-Fuc and L-Gal In contrast, hemagglutination of BGB was not inhibited by any tested monosaccharide, suggesting a strong preference towards this blood group.

Each CRD of IrCLec is predicted to contain a single carbohydrate-binding site, as is typical for C-type lectins, which would theoretically preclude agglutination by isolated domains. This raises the possibility that CRD1 and CRD2 can possess additional, non-canonical binding sites. Those non-canonical carbohydrate-binding sites have been described for at least two C-type lectins (Lefèbre et al., 2024). To our knowledge, no C-type lectin that is able to bind two ligands at once has yet been discovered. Therefore, other explanation is that the recombinant domains are capable of oligomerization. C-type lectin receptors often create either homo- or heterooligomers, for example to improve binding of pathogen glycans or to strengthen signalling (Drickamer and Taylor, 2015).

Glycan array analysis further refined the binding specificity of the individual domains. The results show that all three CRDs of IrCLec bind human blood cell antigens (Figure 8). His-CRD1 and His-CRD2 domains display very similar binding specificity, with the majority of recognized ligands being bound by both. On the other hand, His-CRD3 exhibited a markedly narrower specificity, recognizing only three ligands that were also bound by at least one other domain. All the detected ligands share the common motif Fucα1-2Gal and have complex structure, the only exception being the simple glucose ligand recognized by His-CRD1 and His-CRD3.

**Figure 8.**
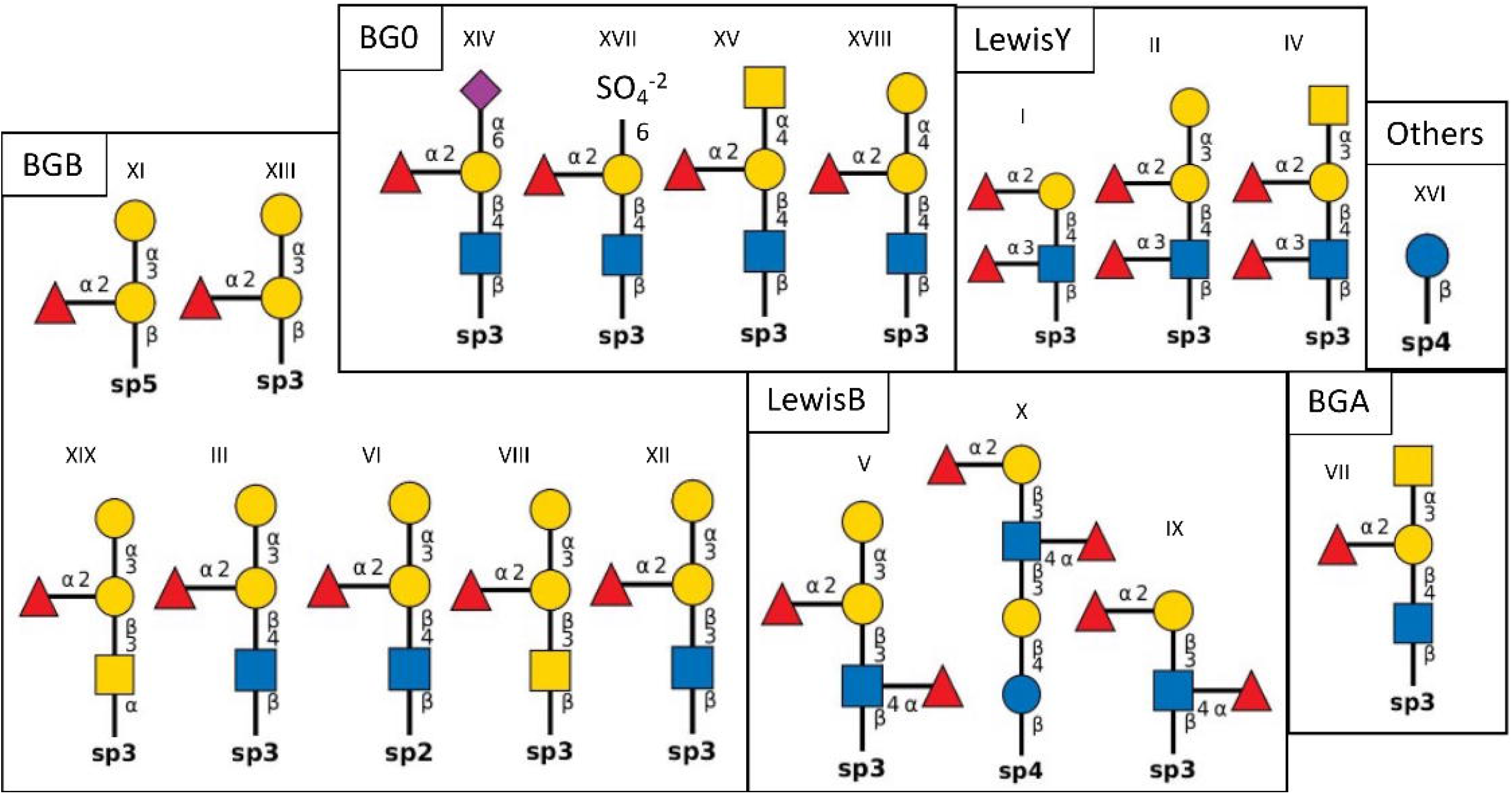
Schematic representation of ligands recognized by all three CRD domains in glycan-array experiments. The ligands are groupped according to a blood group antigen from which they are derived. The symbolic representation of saccharides: yellow circle = D-galactose; yellow square = *N*-acetyl-D-galactosamine; blue circle = D-glucose; blue square = *N*-acetyl-D-glucosamine; red rectangle = L-fucose; purple diamond = *N*-acetylneuraminic acid.

The glucose ligand clearly stands out from the rest of the glycan array results. However, it is not unusual for C-type lectins to recognize glucose as they often have broad specificity, and for example the human mannose receptor is also able to bind this monosaccharide. It was found that the preference of the mannose receptor for glycoconjugates is as follows: L-Fuc = D-Man > D-Glc*N*ac ~ D-Glc > D-Xyl >> D-Gal = L-Ara = D-Fuc, galactose and GalNAc are not recognized (Shepherd et al. 1981). Thus, the fact that none of our CRDs binds mannose while being homologous to MRs is even more interesting finding.

The fine differences in binding selectivity and specificity of the three CRDs can be attributed to the difference in the sequence of their carbohydrate-binding motif. The typical sequential carbohydrate binding motifs of C-type lectins are EPN and WND. These can be found only in the sequence of CRD3. Invertebrate C-type lectins often contain mutations in these motifs, which can lead to alteration of their binding specificity. The EPN motif was previously classified as binding to D-Man, GlcNAc, D-Glc, and L-Fuc (Lee et al., 2011; Zelensky and Gready, 2005). This partially corresponds with the ability of CRD3 to bind L-Fuc and D-Glc, as no interaction with D-Man was observed. In contrast, the QPN motif of CRD1, which was previously described for a lectin from the shrimp *Litopenaeus vannamei* (Wei et al., 2012) and also in *Marsupenaeus japonicus* (Song et al., 2010), was not shown to prefer a particular type of sugars. The QPG motif appearing in CRD2 was identified in a lectin of the snail *Haliotis discus* and showed an affinity for D-Gal, but not to monosaccharides L-Fuc, D-Man, D-Glc, GalNAc, and GlcNAc (Wang et al., 2008). This is inconsistent with the result for CDR2, as only an affinity for L-Fuc was detected. It is therefore clear that predicting the binding specificity is more complicated and considering only a specific sequence motif is not sufficient.

Finally, we predicted the 3D structure of the whole protein as well as separate domains. All three domains retain the typical C-type lectin fold and create three disulfate bond per domain. Single α-helix is present on the C-terminus, with predicted localization within the cell membrane. The predicted spatial arrangement of domains resemble the U-shaped conformation of the mannose receptor (Napper et al., 2001). However, confidence scores for the full-length IrCLec model were lower than those for individual domains, reflecting the current limitations of structure prediction. No arthropod C-type lectin structure has been solved as of now, so comparison of domain arrangement is impossible. Therefore, future structural studies will be essential to confirm the predictions and elucidate the positions of the domains.

## Funding

This work was supported by the Czech Science Foundation (21-29622S). CIISB research infrastructure project LM2018127 funded by the Ministry of Education, Youth and Sport, Czech Republic (MEYS) is gratefully acknowledged for the financial support of the measurements at the CF Biomolecular Interactions and Crystallization and CF Proteomics. JS was supported by Czech Science Foundation (22-25042S). OH was supported by the Czech Science Foundation (no. 25-16064S) and by the grant no. CZ.02.01.01/00/23_020/0008499 funded by the Ministry of Education, Youth and Sport, Czech Republic (MEYS).

## Conflicts of interest

The authors declare no conflict of interest.

## CRediT authorship contribution statement

Kateryna Kotsarenko: Investigation, Writing - Original Draft; Ondřej Hajdušek: Investigation; Martina Rievajová: Formal analysis, Validation, Writing - Review & Editing; Kateřina Bezděková: Methodology; Pavlína Věchtová: Data Curation; Lenka Malinovská: Supervision, Writing - Review & Editing; Eva Paulenová: Investigation; Josef Houser: Investigation; Ján Štěrba: Supervision; Libor Grubhoffer: Supervision Michaela Wimmerová: Conceptualization, Project administration, Funding acquisition.

## Acknowledgments

Core Facility Biomolecular Interactions and Crystallography of CEITEC MU is gratefully acknowledged for the obtaining of the scientific data presented in this paper.

## Notes

### Competing Interest Statement

The authors have declared no competing interest.

