## Supplemental Table S1 for "A newly identified three-domain C-type lectin associated with blood feeding in the tick *Ixodes ricinus*"

Supplementary table 1. List of lectins used for search of the I. ricinus transcriptome

| **Name**  **(or source organism)** | **PDB ID** | **Function** |
| --- | --- | --- |
| Amphibian galectin | 1GAN | Cell adhesion, regulation of growth and apoptosis |
| ERGIC-53 | 1GV9 | Receptor for glycoproteins from endoplasmic reticulum |
| Calnexin | 1JHN | Control of protein folding |
| Mannose phosphate receptor | 1KEO | Biogenesis of lysosomes in higher eukaryotes |
| Lung surfactant protein | 1R13 | Binding to phospholipid membranes of pulmonary alveoli |
| Dectin | 2BPD | Defence against fungal pathogens |
| Malectin | 2JWP | Processing and secretion of glycosylated proteins |
| Fucolectin | 1K12 | Antibacterial protein |
| Tachycitin | 1DQC |  |
| Mannose receptor | 1DQG | Recognition of foreign and self-molecules |
| Congerin | 1C1F | Innate immunity |
| Mannose binding protein | 1MSB |  |
| Tachylectin 5A | 1JC9 |  |
| Tachylectin-2 | 1TL2 |  |
| *Helix pomatia* agglutinin | 2CCV |  |
| Chum salmon lectin CSL3 | 2ZX0 |  |
| Galectin 9 | 3NV1 |  |
| Nematode galectin 9 | 4HL0 |  |
| ZG16p (b-prism) | 3VZF |  |
| DC-SIGN | 3ZHG |  |
| Collectin-K1 | 4YMD |  |
| Selectin | 1G1Q |  |
| H-ficolin | 2J5Z |  |
| FimH | 3ZPD | Adhesin |
| Filamentous hemagglutinin | 1RWR |  |
| Cholera toxin | 1CHP |  |
| Neurotoxin associated hemagglutinin | 1YBI | Hemagglutinin |
| BabA | 5F7L | Adhesion to digestive surfaces |
| SabA | 4O5J |  |
| Salmonella phage P22 | 1CLW | Adhesin |
| *E. coli* phage HK620 | 2VJI |  |
| Morbilivirus | 2ZB6 | Hemagglutinin |
| Murine norovirus | 3LQE | Adhesin, cause of gastroenteritis |
| Bovine-human rotavirus P11 | 4YFW |  |
| Washington university polyomavirus | 3S7X | Adhesin |
| Porcine rotavirus p7 | 3TAY |  |
| Merkel cell polyomavirus | 4FMG |  |
| Bovine coronavirus | 4H14 |  |
| B-lymphotrophic polyomavirus | 4MBX |  |
| *Limax flavus* agglutinin | - | Agglutinin |
| Sialoadhesin | 1URL | Immune system |
| *Limulus polyphemus* – Pentraxin | 1LIM | Innate immunity |
| *Maackia amurensis* agglutinin | - |  |
| *Sambucus nigra* agglutinin II | 3CA4 | Agglutinin |
| Wheat germ agglutinin | 7WGA |  |
| *Maackia amurensis* lectin | 1DBN | Leucoagglutinin |
